# No evidence that polyploids arise during periods of environmental upheaval. A reply to Chen et al. (2026)

**DOI:** 10.64898/2026.07.07.736248

**Authors:** Thomas Marcussen, Andrea S. Meseguer

## Abstract

Ancient whole-genome duplications (WGDs) are thought to have played a major role in plant evolution, but robust inference of the patterns and drivers of polyploid establishment through deep time remains challenging. We re-evaluate the recent large-scale study by Chen et al. (2026), which linked polyploid establishment throughout angiosperm evolution to periods of climatic instability and low species richness. We identify five conceptual and methodological issues that substantially affect these conclusions and collectively undermine the proposed temporal and ecological associations. We hope that clarifying these issues will support future efforts to understand the evolutionary role of polyploidy in plant evolution.

## Main text

Ancient whole-genome duplications (WGDs) are central to understanding plant evolution and diversification. In recent years, the ecological drivers and consequences of polyploidy have attracted considerable attention and have been revisited in numerous large-scale studies and review articles (e.g. Edger et al., 2025; Chen et al., 2026; Sobhanian et al., 2026). In a recent article, Chen et al. (2026) assembled the hitherto largest set of putative ancient WGDs, dated them, and interpreted their temporal distribution as evidence for elevated polyploid establishment during periods of climatic upheaval and declining species richness. Because these conclusions have implications across phylogenetics, paleobiology, and macroevolution, it is important to evaluate the analytical framework on which they rest. Here we highlight five conceptual and methodological issues that have a significant impact on the inferred temporal patterns and ecological interpretations in Chen et al.’s (2026) study: (1) WGD timing is approximated by subgenome divergence time; (2) The timetree is incompatible with the fossil record; (3) The curve-fitting procedure used to infer “polyploid establishment rates” introduces artefactual peaks; (4) The pronounced decline in inferred WGDs toward the present may reflect sampling artefacts; and (5) The species-richness variable is misdefined and its correlation with WGD is algebraic rather than biological. Our aim is to clarify these points constructively and support robust inference about the patterns and drivers of polyploid establishment through deep time.

### Polyploid subgenome divergence time is not WGD time

In their study, Chen et al. (2026) conflated the *divergence time* of the polyploid with the timing of the WGD event itself, and this assumption underlies their inference of the phylogenetic placement and age of candidate WGDs. Although this assumption is frequently made in the literature on polyploidy (e.g., Fawcett et al., 2009; Vanneste et al., 2014; Ren et al., 2018; Cai et al., 2019; Van de Peer et al., 2020; Wu et al., 2020; Guo et al., 2024; Chen et al., 2026), it is also known to introduce significant directional biases in the inferred timing of ancient polyploidy (Doyle & Egan, 2010; Levin, 2013; Dunn & Sethuraman, 2024). By definition, the two events occur close in time in autopolyploids. In contrast, in allopolyploids—which comprise the vast majority of ancient WGDs (>94%) (Dunn & Sethuraman, 2024), including those analysed by Chen et al. — the subgenome divergence time reflects the divergence time of the ancestral diploid species and not the WGD time itself (Doyle & Egan, 2010). The waiting time between these two events can be substantial: for angiosperms in general the waiting time between parental/subgenome divergence time and WGD is expected to be around ∼7 Ma, according to the synthesis presented by Levin (Levin, 2013). He inferred that allopolyploids are most likely to form within the time window between 4–5 Ma and 8–10 Ma after parental divergence (i.e. subgenome divergence), defined by the approximate waiting times for hybrid sterility and cross-incompatibility, respectively. Mathematically, the time interval from subgenome divergence to WGD is a *waiting time* and expected to have an exponential probability density function (Bartoszek et al., 2013; Marcussen et al., 2015), with high variability (mean = standard deviation) and a heavy tail towards high values. In accordance with this, as also noted by Levin (Levin, 2013), polyploid waiting times can occasionally be much longer than the expected ∼7 Ma, and examples not included in his survey comprise ∼16 Ma in the allotetraploid *Viola glabella* (Violaceae) (Marcussen et al., 2011), 17–22 Ma in *Rosa spinosissima* (Rosaceae) (Debray et al., 2021), and 45–50 Ma or more in the recent allopolyploids *Hedlundia hybrida* (Rosaceae) (Liljefors, 1955; Li & Wei, 2025) and *Aesculus carnea* (Sapindaceae) (Upcott, 1936; Xiang et al., 1998) —all of these belong in lineages which have a rich fossil record (Manchester, 2001; DeVore & Pigg, 2007; DeVore & Pigg, 2016; Marcussen et al., 2022) and have been reliably dated. In summary, Chen et al. dated the wrong biological event, and in doing so introduced a directional temporal bias in their estimated “WGD” times. Because this approximation enters at the earliest stage of the Chen et al. pipeline, increasing the WGD age estimates by possibly ∼7 Ma on average, it affects all downstream analyses and putative correlations with climatic events.

### The timetree is in conflict with the fossil record

The chronogram obtained by Chen et al. (2026) (https://doi.org/10.5281/zenodo.19224723; file: Maintree.txt) is incompatible with widely accepted minimum ages for numerous major angiosperm clades. These minima are well established in the paleobotanical literature (e.g. Friis et al., 2011; Iles et al., 2015; Manchester et al., 2015) and based for the most part on macrofossils with numerous diagnostic characters whose placements within the indicated clades are not considered controversial. Although Chen et al. included 44 deep fossil calibrations in their study, these were typically placed at the stem node of large, inclusive clades (often entire orders) disregarding any information about more nested placements of the fossils within the lineage, leading to systematic underestimation of ages at middle and shallow phylogenetic depths. For example, the chronogram of Chen et al. places crown Juglandaceae *sensu stricto* at 17.9 Ma despite unequivocal juglandaceous fossils from ∼65 Ma onwards (Manchester, 1987; Lyson et al., 2019); crown Maleae (Rosaceae) at 22.8 Ma, *Fraxinus* (Oleaceae) at 21.2 Ma, and *Physalis–Capsicum* (Solanaceae) at 18.8 Ma despite abundant middle Eocene (∼50 Ma) records of these lineages (DeVore & Pigg, 2007; Mathewes et al., 2021; Deanna et al., 2023); and *Populus* (Salicaceae) at 25.0 Ma despite having a fossil record from 56 Ma onwards (Manchester et al., 1986). Similar discrepancies occur across the phylogeny, e.g. for *Aesculus* (Sapindaceae) which has an estimated age at 42.8 Ma but a considerably older fossil record (Manchester, 2001), for Araliaceae estimated at 56.0 Ma (Friis et al., 2011; Manchester et al., 2015), Arecales crown 51.9 Ma (Iles et al., 2015), Malpighiales crown 77.0 Ma (Friis et al., 2011), Malvaceae crown 41.8 Ma (Friis et al., 2011), Myrtales crown 75.2 Ma (Friis et al., 2011), Nymphaeales crown 71.4 Ma (Friis et al., 2009), Poaceae crown 71.8 Ma (Wu et al., 2017), *Prunus* 40.3 Ma (Benedict et al., 2011), Rhamnaceae (per *Ziziphus*) 55.2 Ma (Friis et al., 2011), and *Rhododendron* (Ericaceae) 34.8 Ma (Collinson & Crane, 1978). Conversely, *Tetracentron–Trochodendron* (Trochodendraceae) is placed at 108.2 Ma despite neither lineage having a fossil record older than Paleocene (Manchester et al., 2021). It is not our purpose to discuss the fossil record of each of these lineages, but merely to show that the age discrepancies in Chen et al.’s data are numerous, often exceeding 20–40 Ma, and that they affect the entire phylogeny, and therefore downstream analyses based on these dated WGD events. Chen et al. do not evaluate these inconsistencies, yet their conclusions rely directly on absolute divergence and WGD timing.

### The reported peaks in WGD establishment rates are analysis artifacts

The curve-fitting procedure applied by Chen et al. (2026) imposes temporal structure that is supported neither by the dated phylogeny itself nor by the dated subgenome divergence events– in this paragraph, we refer to the latter as “WGD” events to remain consistent with the terminology adopted by Chen et al. (but see the previous paragraph). Both come with substantial age uncertainties, on average 24.8 Ma for the “WGDs” (90% highest-credibility region (HCR); minimum 5.3 Ma; maximum 62.0 Ma; Table S11) and 27.5 Ma for the node heights in the dated phylogeny (95% highest probability density (HPD); minimum 0.07 Ma; maximum 81.6 Ma; https://doi.org/10.5281/zenodo.19224723, file “Maintree.txt”). However, both sources of uncertainty are ignored in the analytic pipeline of Chen et al.: they first collapsed the posterior “WGD” age distributions to single point estimates and binned them into 1 Ma intervals. From this point-binned series, Chen et al. compute a “WGD establishment rate” defined as the number of WGDs per lineage per Ma, thereby neglecting node-age uncertainty in the timetree as well. They then apply a narrow 4 Ma sliding window to estimate a continuous temporal rate of WGD establishment. Crucially, the selected window size of 4 Myr is less than one sixth of the average uncertainty reported for both “WGDs” and node heights (and is narrower than the HCR for any “WGD”; minimum 5.3 Ma). Because age uncertainties are discarded in the first step of the analysis, applying such a narrow sliding window size inevitably introduces artificial temporal structure at a finer scale than the resolution supported by the underlying posterior distributions. As a result, any apparent peaks or troughs at this scale are likely artefacts of the binning and smoothing procedure rather than features of the dated events themselves.

To distinguish structure present in the data from structure introduced by the analytical approach of Chen et al. (2026), we re-analysed their data by estimating the number of “WGDs” (in reality subgenome divergence events) per unit time and the number of species per unit time (Figure 1A), as well as their ratio (Figure 1B). This information was acquired directly from the age distributions provided by the authors, without binning, sliding windows, or LOESS smoothing, and preserves the original age uncertainty in the data (Figure 1; Methods). The R script covering all the steps is available as a supplementary file. The unsmoothed subgenome divergence to species ratio curve (Figure 1B) shows a broad maximum around 90–50 Ma and a decrease to the present, and only four of the nine peaks inferred by Chen et al. are recovered, but only as minor features of the graph. Our failure to recover the narrow, pronounced peaks reported by Chen et al. (Figure 3A in Chen et al.), demonstrates that those features arose from their smoothing pipeline rather than from the underlying age posteriors. Because the peaks were used by Chen et al. for downstream significance tests and correlations, the corresponding inferences are not supported.

**Figure 1.**
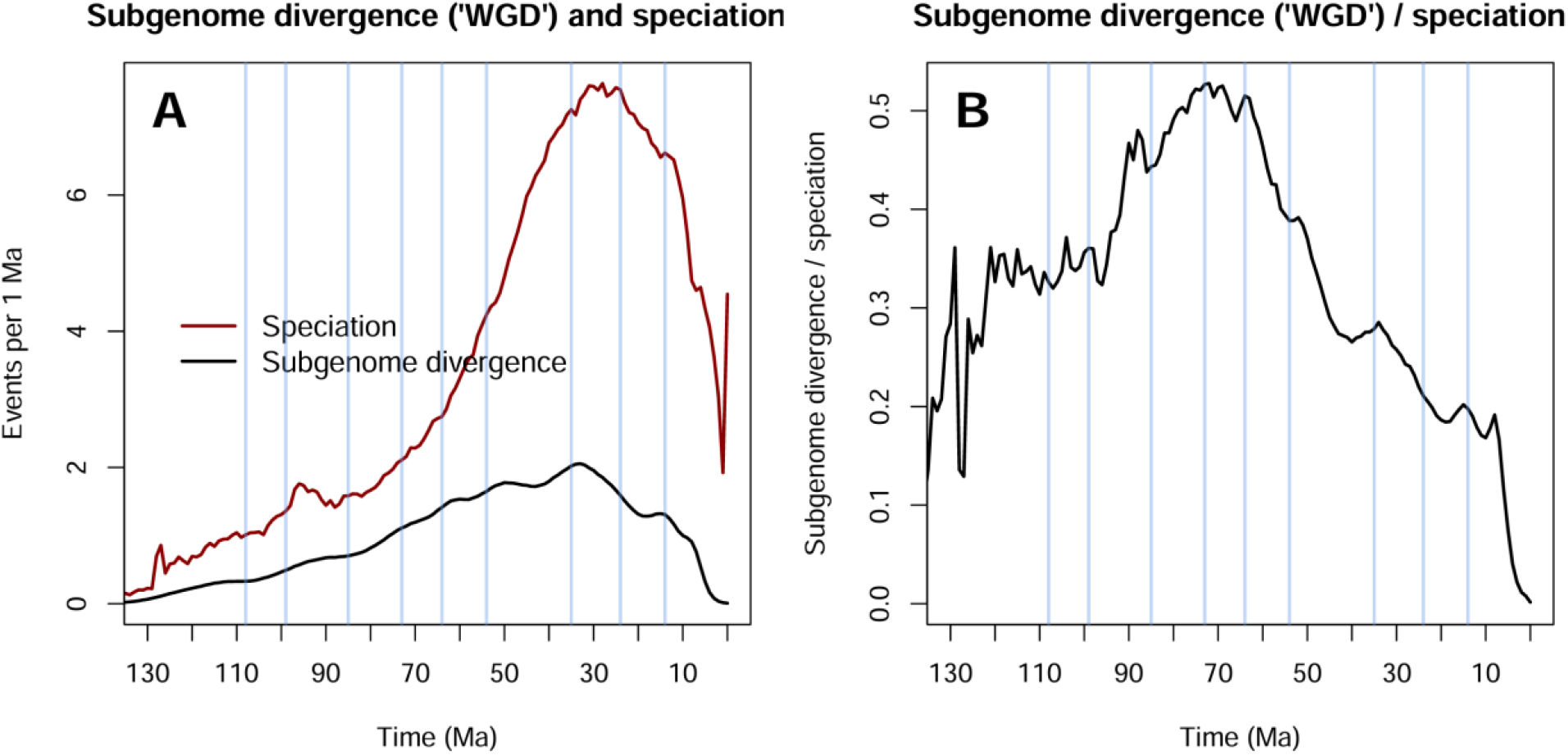
Reconstructed “WGD” curves from Chen et al. (Chen et al., 2026) data. (**A**) Expected number of subgenome divergence events (treated as “WGD” events by Chen et al.; black line) and speciation events (dark red line) per 1 Ma time bin, obtained by sampling 100,000 ages from the reported 90% posterior interval of each WGD, and 100,000 ages for the 95% HPD interval of each internal node in the timetree. Vertical blue lines mark the nine peaks in “Excess WGD establishment rate” inferred by Chen et al. The abrupt increase of the speciation curve near 0 Ma reflects the sampling of several taxon pairs with recent divergence times. (**B**) Ratio of subgenome divergence events to homoploid speciation events per 1 Ma time bin, derived directly from the curves in panel A. No smoothing, sliding windows, or curve-fitting procedures were applied; curves reflect the posterior age distributions and HPD-based lineage-through-time structure of the original data.

### The collapse in WGD toward the present is most likely a sampling artifact

Any interpretation of the occurrence of polyploidisation through time ultimately rests on the representative and correct detection of these events through time. However, both the original WGD establishment rate curve reported by Chen et al. ((2026) Figure 3B) and our re-analysed “WGD”/speciation curve (Figure 1B) show strong declines to zero towards the present. This pattern stands in stark contrast to the known high incidence of polyploidisation in recent times, where it is estimated that up to ∼30% of angiosperm speciation events involve polyploidy (Mayrose et al., 2011) and that 35% of extant vascular species are within-genus polyploids (Wood et al., 2009). Indeed, among the species selected by Chen et al., we could demonstrate that at least 20 uncontested cases of young allopolyploids had been overlooked by Chen et al.: i.e. *Arachis hypogaea* (tetraploid; 4x) (Zhuang et al., 2019), *Avena sativa* (6x) (Liu et al., 2017), *Brassica napus* (4x) (Chalhoub et al., 2014), *Camelina sativa* (6x) (Kagale et al., 2014), *Chenopodium quinoa* (4x) (Zou et al., 2017), *Cuscuta campestris* (4x) (Cerda-Herrera et al., 2025), *Eragrostis tef* (4x) (VanBuren et al., 2020), *Gossypium barbadense* and *G. hirsutum* (4x) (Senchina et al., 2003), *Nicotiana tabacum* (4x) (Leitch et al., 2008), *Panax ginseng* (Song et al., 2024) and *P. quinquefolius* (4x) (Lei et al., 2025), *Perilla frutescens* (4x) (Tamura et al., 2023), *Pogostemon cablin* (4x) (Shen et al., 2022), *Rhynchospora pubera* (8x) (Hofstatter et al., 2022), *Thinopyrum intermedium* (6x) (Sun et al., 2025), *Thymus quinquecostatus* (4x) (Sun et al., 2022), *Triticum aestivum* (6x) and *T. turgidum* (4x) (Marcussen et al., 2014), *Vaccinium corymbosum* (4x) (Colle et al., 2019)). As the subgenomes associated with these WGDs all diverged within the last 0–35 Ma (according to the dates obtained by Chen et al.) these events would have contributed to “filling the gap” towards the present in their data. Therefore, the pronounced decline in both inferred “WGDs” and speciation from ∼30 Ma to the present (Figure 1A) is demonstrably caused by the under detection of recent WGD events, rather than reflecting a biological signal, which affects the validity of their downstream inferences about diversity or environmental drivers.

### The association between WGD and species richness is a misinterpretation

That WGD establishment is negatively correlated with species richness, i.e. the number of species per defined geographic unit, is well established in polyploid research (Rice et al., 2019). However, the inference of this relationship also with the current dataset rests on a statistical relationship that is algebraically predetermined and conceptually mischaracterized in the text of Chen et al. (2026). The authors define the WGD establishment rate as the number of WGDs per lineage per unit time, and then regress this rate against lineage counts per unit time. Because lineage count is embedded in the nominator and in the denominator of the response variable, any negative association between WGD establishment rate and lineage count is tautological, and at least in part, mathematically expected from the definition of the statistic. It therefore does not, by itself, constitute an empirical finding of diversity dependence.

Moreover, Chen et al. (2026) misinterpret this algebraic relationship as evidence that WGD establishment is higher when “species richness” is lower, treating lineage count and species richness as interchangeable descriptors. However, lineage-through-time (LTT) counts inferred from phylogenies merely reflect the accumulation of surviving and sampled lineages within the dataset rather than the true global species richness through time – extinct lineages that left no descendants and non-sampled species are absent from the reconstructed tree (Nee et al., 1994) and thus from the species counts. Under substantial extinction, LTT curves may severely underestimate past diversity, and distinct diversification scenarios —including “stasis and rate-shift diversification” dynamics, waxing-and-waning, diversity decline, or mass extinction events—can generate similar LTT patterns (Sanmartin & Meseguer, 2016; Louca & Pennell, 2020), limiting the utility of LTT curves alone as direct evidence of historical diversity. The regression they present therefore tests a mathematical identity rather than a biological hypothesis, and the subsequent ecological interpretation— that polyploid establishment is suppressed under species-rich conditions—does not follow from the data or from the quantities actually analyzed.

## Conclusion

Taken together, these five points indicate that the temporal clustering and ecological correlations reported by Chen et al. (2026) cannot yet be interpreted as evidence for elevated polyploid establishment during periods of environmental upheaval and low species richness. We emphasise that our critique concerns the methodological implementation rather than the broader hypotheses motivating the study, which we regard as timely and worthy of further investigation. We hope that clarifying these methodological issues will support future efforts to understand the role of polyploidy in plant evolution through deep time.

## Methods

To reconstruct the temporal distribution of the subgenome-divergence times reported by Chen et al. (Chen et al., 2026) (treated as “WGD” times in their study), we extracted the WGD identifiers, consensus ages, and 90% highest-confidence ranges (HCRs) from *Supplementary Table 4*. For each WGD, we fitted a normal distribution whose central 90% interval matched the reported HCR and drew 100,000 ages from this distribution. Internal node ages in the timetree were obtained by extracting the 95% HPD bounds embedded in the Newick annotations of *Maintree*.*txt*; for each node, we assumed a uniform distribution across its HPD interval and drew 100,000 ages. All sampled ages were assigned to 1 Ma bins to obtain the expected number of WGD events, homoploid speciation events, and their ratio through time (Figure 1). This procedure yields fully unsmoothed pWGD(t) curves that reflect the posterior age distributions and HPD-based lineage-through-time structure directly, without any sliding windows, LOESS smoothing, or curve-fitting.

